# Emergence and interstate spread of highly pathogenic avian influenza A(H5N1) in dairy cattle

**DOI:** 10.1101/2024.05.01.591751

**Authors:** Thao-Quyen Nguyen, Carl Hutter, Alexey Markin, Megan Thomas, Kristina Lantz, Mary Lea Killian, Garrett M. Janzen, Sriram Vijendran, Sanket Wagle, Blake Inderski, Drew R. Magstadt, Ganwu Li, Diego G. Diel, Elisha Anna Frye, Kiril M. Dimitrov, Amy K. Swinford, Alexis C. Thompson, Kevin R. Snevik, David L. Suarez, Erica Spackman, Steven M. Lakin, Sara C. Ahola, Kammy R. Johnson, Amy L. Baker, Suelee Robbe-Austerman, Mia Kim Torchetti, Tavis K. Anderson

## Abstract

Highly pathogenic avian influenza (HPAI) viruses cross species barriers and have the potential to cause pandemics. In North America, HPAI A(H5N1) viruses related to the goose/Guangdong 2.3.4.4b hemagglutinin phylogenetic clade have infected wild birds, poultry, and mammals. Our genomic analysis and epidemiological investigation showed that a reassortment event in wild bird populations preceded a single wild bird-to-cattle transmission episode. The movement of asymptomatic cattle has likely played a role in the spread of HPAI within the United States dairy herd. Some molecular markers in virus populations were detected at low frequency that may lead to changes in transmission efficiency and phenotype after evolution in dairy cattle. Continued transmission of H5N1 HPAI within dairy cattle increases the risk for infection and subsequent spread of the virus to human populations.

## Introduction

Highly pathogenic avian influenza (HPAI) viruses have critical consequences for animal health, the agricultural economy, and may have pandemic potential. HPAI related to the goose/Guangdong 2.3.4.4 hemagglutinin (HA) H5NX phylogenetic clade has spread to nearly 100 countries (*1*), infections resulted in mortality events, and it is recognized as a panzootic, crossing multiple species barriers. Additionally, HPAI virus circulation is enzootic in Europe, with consistent detections representing a potential shift in the biology and transmission of HPAI (*2*). Following an initial trans-Atlantic incursion in late 2021 (*3, 4*), the HPAI H5N1 clade 2.3.4.4b virus caused widespread outbreaks across North America (*5*). The outbreaks resulted in extensive mortality events in wild bird species, mortality and culling of poultry when detected in agricultural systems, a significant number of interspecies transmission events into wild mammals, and human infections.

The relatively frequent and recent transmission of HPAI clade 2.3.4.4 between avian species and mammals globally has resulted in persistence of genetic features associated with mammalian adaptation. Experimental studies on a subset of viruses from the HPAI clade 2.3.4.4 have demonstrated that they can bind to both human α2,6-linked and avian α2,3-linked sialic acid receptors (*6, 7*). Genomic analyses have documented that approximately half of the sequences from mammals globally within the HPAI H5N1 clade 2.3.4.4b have amino acid signatures in the polymerase basic (PB) 2 protein that have been associated with mammalian adaptation through enhanced viral replication, host-specific polymerase activity, and temperature sensitivity (E627K, D701N, and/or T271A) (*5, 8*). Additionally, the introduction of HPAI H5N1 into farmed mink in Europe in 2022 provided evidence that transmission to, and within, a population of mammalian hosts could result in mutations in the hemagglutinin associated with human receptor recognition and in the neuraminidase protein that affected sialic acid binding in a manner similar to human influenza A viruses (*9–11*). These results are particularly important as from January 2022 to April 1, 2024 there were 13 reported human cases of H5N1 from the HPAI clade 2.3.4.4b worldwide, with some having severe consequences, including mortality (*12*). Consequently, it is critical to determine how evolution of the HPAI clade 2.3.4.4b in wild birds and the associated spillovers and transmission in mammals impacts genomic and phenotype features that alter the potential for human infection and transmission (*13, 14*).

On March 25 2024, HPAI H5N1 clade 2.3.4.4b was confirmed in dairy cattle in Texas following multistate reports of decreased milk yields. Shortly thereafter, the virus was identified in cattle in eight other US states by members of the National Animal Health Laboratory Network (NAHLN) (*15–17*). Virus was predominantly found in mammary tissue and milk. It was also detected in cats and peridomestic animals that died on affected premises. Overall, the detection of influenza A virus (IAV) in cattle has been rarely documented (*18, 19*), but there is prior evidence for its replication within the mammary gland (*20, 21*), association with a reduction in milk yield (*22*), and experimental studies have demonstrated that bovine calves are susceptible to infection and may asymptomatically shed virus (*23*). The goal of this study was to analyze available genetic sequence data collected following the introduction of HPAI H5N1 in late 2021 into the Atlantic flyway of North America and its onward circulation and reassortment with North American wild-bird origin low pathogenicity viruses that resulted in over 100 distinct genotypes (*24*). These data were synthesized with newly generated whole-genome sequence data and epidemiological information from the outbreak among US dairy cattle to understand when the interspecies transmission event occurred and the consequences of animal movement on the persistence and evolution of the virus. To achieve this, we performed phylodynamic analysis of the US HPAI H5N1 viruses detected in dairy cattle alongside epidemiologically linked wild bird, poultry, and peridomestic animal data. In addition, we present a within-host evolutionary assessment of the virus to determine how onward transmission in dairy cattle affects genomic diversity and whether this increases the potential for this host to serve as a reservoir for IAV with zoonotic potential.

## Results

### H5 clade 2.3.4.4b introduction into the United States

The H5N1 clade 2.3.4.4b genotype A1 (*24*) was first identified in wild birds collected December 2021 (*3*). Over 100 genotypes representing different gene constellations have been characterized but 70% of all US viruses fall into only 7 genotypes (*25*). Despite these genetic differences, spillover events have been associated with the predominance of a genotype rather than any specific link between genotype and host specificity. Introduction of genotype A3 was identified in April 2022 likely via the Pacific flyway followed by A4. Another two introductions were identified via the Atlantic flyway, including an A6 Eurasian virus that had a reassorted Eurasian neuraminidase (H5N5). Widespread detections in wild bird populations (*26*) continue to result in point source spillovers to poultry (*27*) but there is limited evidence for lateral transmission among poultry flocks. Virus of multiple genotypes have also been detected in approximately 20 mammal species in the US often related to mortality or severe neurologic signs, but these appear to be dead end hosts (*28, 29*). In early March 2024, HPAI H5N1 was identified in neurologic goat kids on a farm where poultry had recently been depopulated for HPAI; this event was unrelated to the dairy event and involved a different virus genotype.

### Epidemiological investigations detected HPAI H5N1 in dairy cattle across the United States

In late January 2024, production veterinarians observed dairy cattle displaying unexplained reductions in milk production, decreased feed intake, and changes in milk quality. Members of the National Animal Health Laboratory Network (NAHLN) identified influenza A virus in milk and a few nasal swabs from a Texas dairy and forwarded samples to the National Veterinary Services Laboratories (NVSL) for confirmatory testing as epidemiologic investigations continued. Testing revealed the presence of H5N1 clade 2.3.4.4b genotype B3.13. Shortly after the identification of HPAI H5N1 genotype B3.13 in Texas, it was confirmed in additional Texas herds and herds in other states (*16*). Samples collected between 7 March 2024 and 8 April 2024 have virus characterized as genotype B3.13 from 26 dairy cattle premises across eight states and six poultry premises in three states (Data S1). The NVSL conducted whole genome sequencing and analysis using a custom software program that identified transmission chains based on genomic similarity (vSNP) to provide rapid feedback in support of epidemiologic investigations (Data S2, Fig. S1). The phylogenetic and available epidemiologic data indicated that the genotype B3.13 virus was being moved between dairy cattle premises, as well as domestic poultry premises, via multiple transmission routes (Fig. 1). Detections of the B3.13 genotype in cattle locations that have no known epidemiologic links to confirmed premises suggest there are affected herds that have not yet been identified.

**Fig. 1.**
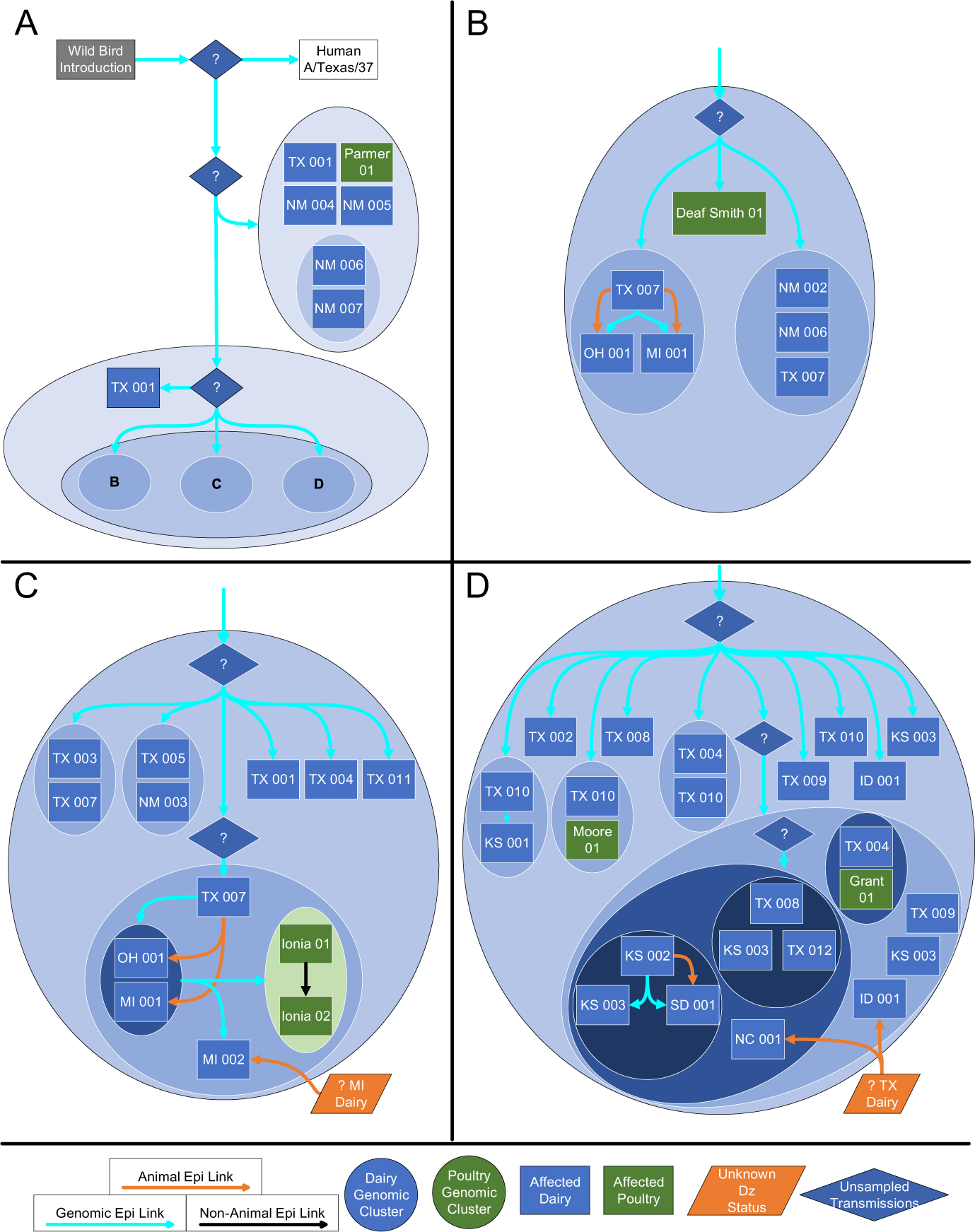
Putative transmission pathways of HPAI H5N1 clade 2.3.4.4b genotype B3.13 supported by epidemiological links, animal movements, and genomic analysis. Three lines of evidence support a single interspecies transmission event followed by lateral spread within dairy cattle with onward transmission to poultry. This visualization represents 26 dairy cattle premises (blue boxes) and six poultry premises (green boxes). Herds with unknown disease status are colored orange, and putative unsampled transmission links are indicated with diamonds. Genomic clusters, specifically those strains that are more similar to each other than other clusters, are bounded by progressively smaller circles. Genomic links are light blue arrows, confirmed animal movements are orange arrows, and non-animal links are black arrows. Premises that appear more than once indicate the presence of genetically distinct viruses sequenced from independent samples on those premises. (**A**) The gray box represents an unsampled wild bird-origin common ancestor. The white box is the B3.13 genotype human detection similar but distinct from the sampled cattle and poultry clusters. Two major groups emerged from an unsampled common ancestor: one containing dairy premises in Texas and New Mexico along with Parmer 01 poultry, and another containing TX 001 and three clusters labeled (**B**), (**C**), and (**D**) respectively. An unsampled common source links the premises shown in B, C, and D within each of the clusters. (**B**) Two known animal movements occurred in this cluster: TX 007 to OH 001, and TX 007 to MI 001. These are shown in panel C, indicating that simultaneous transmission of genetically distinct viruses occurred. (**C**) MI 001 was genetically and temporally linked to the Ionia 01 poultry premises, and Ionia 01 was genetically, epidemiologically, and temporally linked to Ionia 02, suggesting a lateral transmission chain. Prior to detection, MI 002 received an animal movement from an unknown Michigan dairy premise of unknown disease status. (**D**) Phylogenetic analysis suggested several directional transmissions: TX 010 to KS 001, KS 002 to KS 003, and KS 002 to SD 001. The link from KS 002 to SD 001 was supported by an animal movement. Prior to detection, NC 001 and ID 001 each received an animal movement from the same Texas dairy of unknown disease status.

### Genomic epidemiology demonstrated a single interspecies transmission event

To determine when HPAI H5 clade 2.3.4.4b virus was introduced into cattle in the US, we conducted a phylogenetic analysis using whole genome sequence data collected from poultry, wild birds, and mammals (Fig. 2, Fig. S2). From 2022 to present, clade 2.3.4.4b viruses have been reported in over 9,000 wild birds in at least 163 species across 49 states and Washington D.C., in over 200 mammals in at least 20 species across 29 states and Washington D.C., and over 1,120 poultry flocks across 48 states. For the HA gene segment, the cattle isolated H5N1 2.3.4.4b sequences clustered within a single monophyletic clade strongly supporting a hypothesis for a single spillover event followed by lateral transmission. The single spillover was also supported by the phylogeny reconstructed from concatenated whole genome sequences of B3.13 genotype strains (Fig. 2), and the single introduction pattern was congruent across the PB2, PA, NP, NA, MP, and NS gene segments (Fig. S3-S6). For PB1, over 97% of cattle sequences also formed a single monophyletic clade; the remaining sequences were placed in a parental clade due to high similarity among the gene sequences and low phylogenetic signal (Fig. S6). The single introduction hypothesis was further supported by single nucleotide variant calling analyses (Data S2, Fig. S1). The inferred evolutionary rate was 6.23×10^−3^ substitutions/site/year (95% highest posterior density (HPD), 5.29×10^−3^-7.19×10^−3^). The time to the most recent common ancestor (TMRCA) for the HA segment for the cattle clade 2.3.4.4b HPAI H5N1 sequences was estimated as December 9, 2023 (95% HPD, October 12, 2023, to January 26, 2024. A clade 2.3.4.4b genotype B3.13 infection in a dairy worker was diagnosed at the end of March 2024 and was similar to the HPAIV from cattle. The TMRCA for the cattle group and the human virus was estimated as November 10, 2023 (95% HPD, September 2, 2023, to December 24, 2023). These two groups shared a TMRCA that was estimated as October 19, 2023 (95% HPD, August 22, 2023, to December 9, 2023). More accurate TMRCA estimates may be inferred using host-specific molecular clocks models (*30*), but our dates were supported by TMRCA estimated using maximum likelihood methods (Fig. 2) and they are feasible given associated epidemiological information (Fig. 1). These data support a single introduction event from wild bird origin virus into cattle, likely followed by limited local circulation for approximately 4 months prior to confirmation by USDA.

**Fig. 2.**
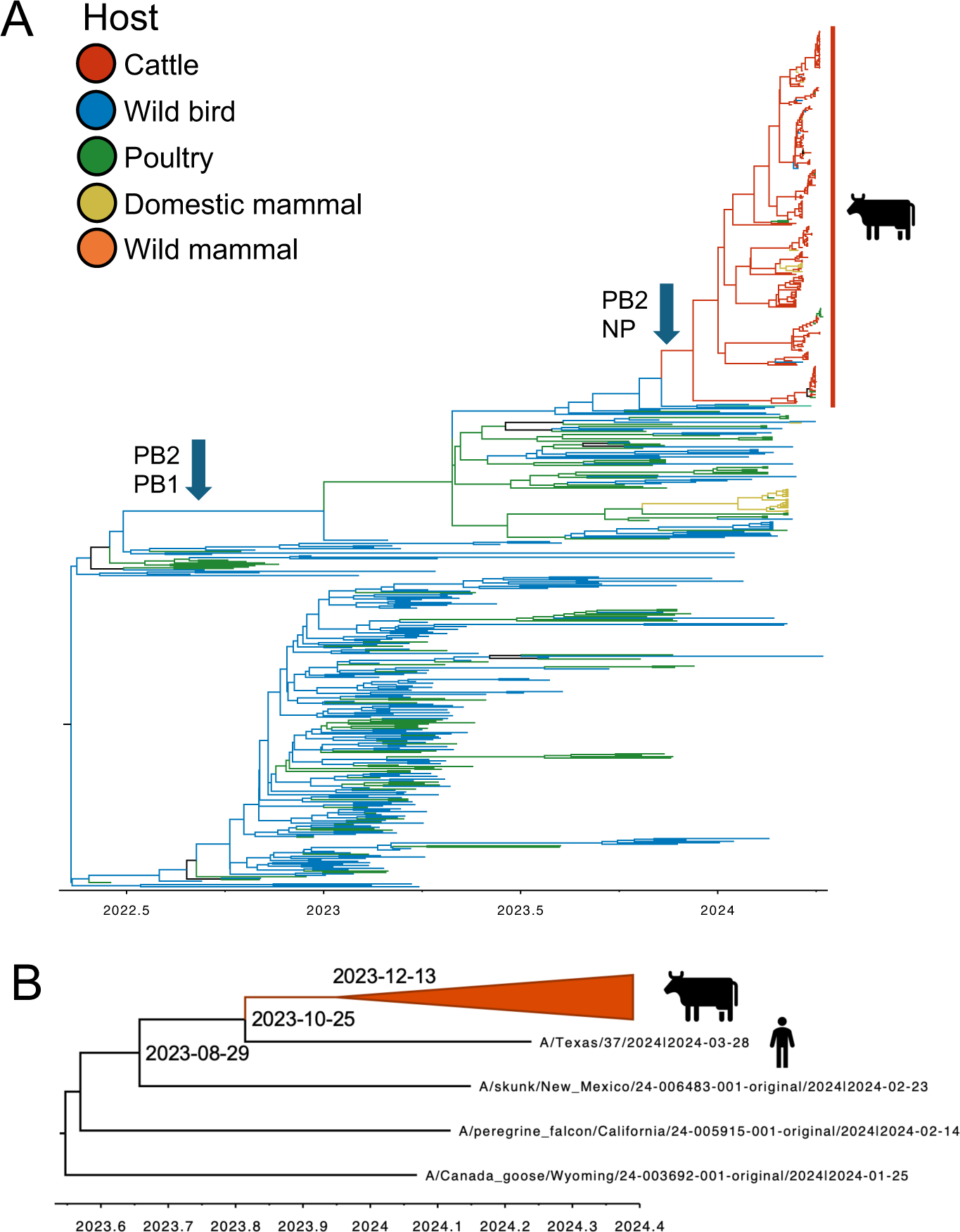
Evolutionary history of H5N1 2.3.4.4b in North America prior to and during the introduction and emergence in US cattle. (**A**) A time-scaled phylogeny of the HA gene between December 2021 and April 2024 demonstrating the single introduction of the virus from wild birds to dairy cattle with an estimated date of December 2023 (95% credible interval: October 2023 – January 2024). B3.13 strains inherited PB1, PA, HA, NA, MP, and NS genes from a B3.6 ancestor and acquired different PB2 and NP genes from North American LPAI viruses. This phylogeny was paraphyletically subsampled to maintain tree topology and demonstrate the 12 subsequent spillovers from cattle to domestic cats, poultry, and peridomestic animals and is presented in the supplemental figures. (**B**) A simplified visualization of a hypothesis on the evolutionary history of the H5N1 2.3.4.4b B3.13 genotype that emerged in cattle and a range of other mammalian and avian hosts. This time-scaled phylogeny was inferred using maximum likelihood methods from concatenated genomes: inferred dates presented at nodes for the most recent common ancestor were congruent with those estimated using Bayesian methods.

### Transmission within cattle resulted in subsequent interspecies transmission events to domestic cats, poultry, and peridomestic animals

The phylogenetic tree topology indicated that following the introduction, the virus persisted in cattle populations, with subsequent evidence for transmission from cattle into poultry and peridomestic animal species (Fig. 1-2, Figs. S2, S7). There were as many as five cattle to poultry, one cattle to raccoon, two cattle to domestic cats, and three cattle to wild bird transmission events. These were further supported by epidemiological information, as these animals were collected from premises with cattle where genotype B3.13 HPAIV was identified. Our Bayesian discrete state analysis (Fig. 3) that quantified the movement of HPAIV between six different host categories (poultry, wild bird, cattle, wild mammal, domestic cat, and humans) demonstrated sufficient evidence to support the proposition of HPAI in cattle resulted in infections in other hosts. We cannot exclude the possibility that this genotype is circulating in unsampled locations and hosts as the existing analysis suggests that data are missing and undersurveillance may obscure transmission inferred using phylogenetic methods (*31*). The gap in data is highlighted by the human infection with genotype B3.13 HPAIV where the HA gene sequence was not nested within cattle HA gene sequences. This could indicate that HPAIV in unsampled cows were the source of infection or within-host evolution resulted in divergence sufficient to result in a different phylogenetic grouping. It is most likely, however, that asymptomatic transmission and undersurveillance in epidemiologically important populations drove this pattern. Our analysis of transmission chains within the cattle B3.13 clade using a phylogenomic approach suggested unsampled transmission in late 2023 and early 2024 (Fig. S8), and the TMRCA indicates there may have been 4 months of circulation prior to confirmation by USDA. However, given the decline in milk production in highly monitored dairy herds, it is unlikely that the spillover occurred significantly outside of the described TMRCA ranges.

**Fig. 3.**
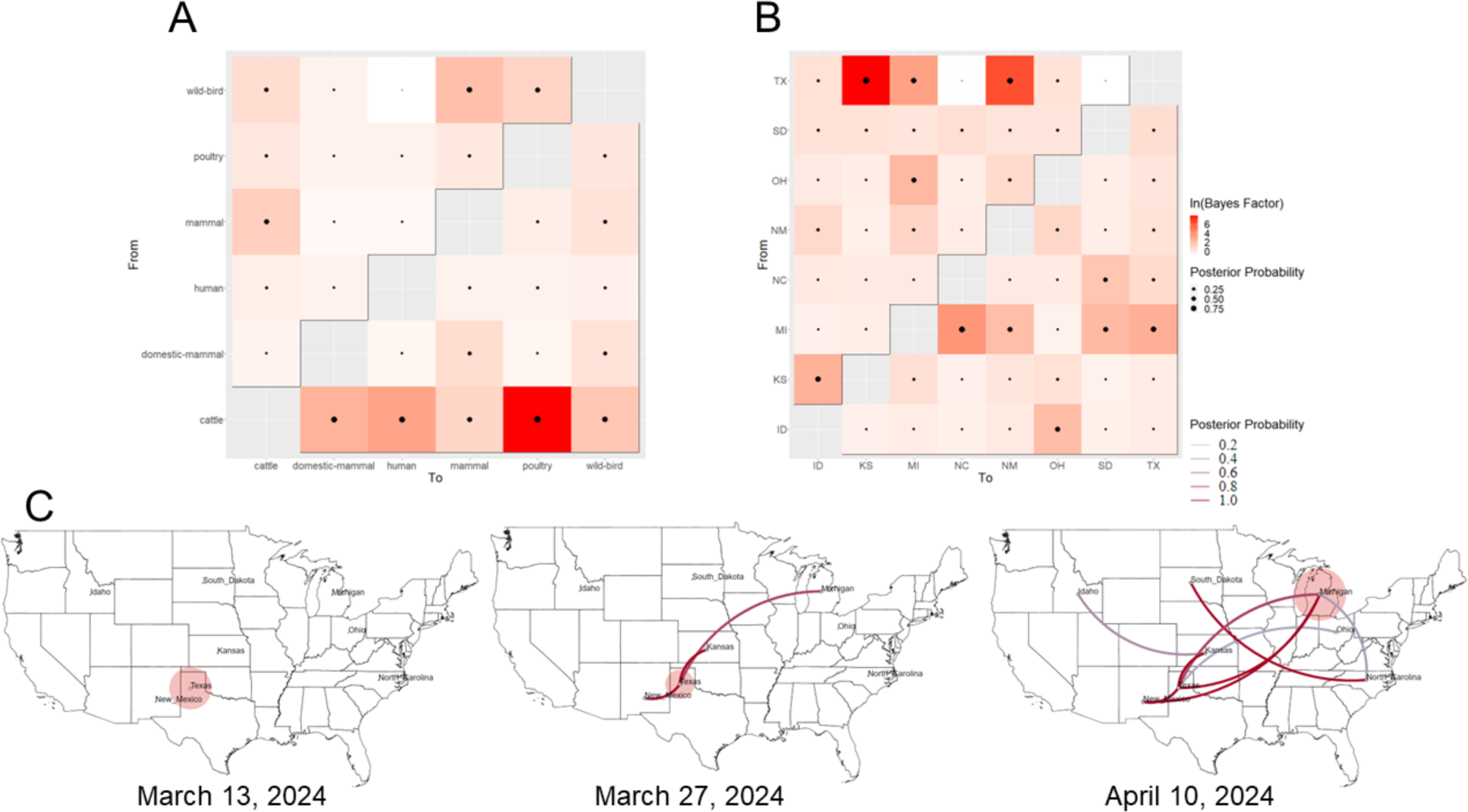
Bayes factors for inferred movement between different discrete traits of H5N1 clade 2.3.4.4b viruses demonstrating the frequency of movement. Bayes factors and posterior probabilities were plotted to demonstrate the frequency and support for the movement of the clade 2.3.4.4b B3.13 HA gene between different host species (**A**) or US locations with documented detections of 2.3.4.4b (**B**) with the source host species or US state plotted on the *y*-axis, and the recipient host species or US state plotted on the *x*-axis. This analysis was conducted on the subsampled and aligned hemagglutinin dataset (A), and an extracted subset of the data that covered the monophyletic clade of cattle HA clade 2.3.4.4b genes (B). When imposing a Bayes factor > 3 with an associated posterior probability greater than 75%, there were supported transitions from Texas to Kansas, New Mexico, and Michigan, and from Michigan to North Carolina. (**C**) Transitions between locations were inferred using a discrete trait model across time and using 14-day timepoints of the spatial spread corresponding to beginning, middle, and end of our dataset. The centroid of each state was used: for Texas we located the point in the panhandle where dairy cattle are commonly farmed. The circular polygon reflects the number of lineages within a location and eastward movements are depicted by lines with a downward curve, while westward movements are depicted by lines with an upward curve.

### Reassortment in migratory bird populations resulted in a new genotype associated with emergence in cattle

From January 2022 to April 2024, more than 482 commercial and 645 backyard flocks tested positive for HPAI (*5, 27*). A major component in the dissemination of the 2.3.4.4b viruses is movement across four migratory flyways, and infection of nearly 200 wild avian species (*5*). Extensive genetic reassortment with existing North American wild bird LPAIVs is a consequence of the host- and geographic-breadth. This generated a spectrum of different genotypes, with the HA, NA, and M genes demonstrating preferential pairings similar to mammalian adapted IAV (*32*) and transient detections of reassorted viruses with different PB2, PB1, PA, NP, and NS genes (*5*). In this study, we detected 243 putative reassortment events across the HA phylogeny (Fig. S10). These events were associated with PB2 (137/243), PB1 (78/243), PA (36/243), NP (126/243), and NS (52/243) gene segments. Most of these events did not persist. However, we identified 24 major reassortment events that resulted in new genotypes with more than 20 downstream detections. The spillover into cattle was preceded by a reassortment event(s) involving different PB2 and NP genes, likely derived from wild bird LPAI in late 2023. This reassortment resulted in the B3.13 genotype that is maintained across the clade of epidemiologically linked cattle samples with no evidence for further reassortment following the spillover. The NP gene acquired by the reassortment event may have resulted in a phenotype change that mediated the emergence of this virus genotype in cattle. The NP gene is associated with multiple processes in the virus lifecycle (*33*) and was implicated in increasing the transmission efficiency of IAV in the swine host (*34*).

### HPAI H5N1 dispersed across the United States tracking cattle movement

Epidemiological records documented dairy cattle movement from a Texas herd (at shipment, HPAI status was unknown) to North Carolina and Idaho (Fig. 1). Records also indicated that asymptomatic dairy cattle, some that were subsequently diagnosed as HPAIV positive, were moved from a Texas herd to Michigan and to Ohio, and a Kansas herd to South Dakota. We also applied an asymmetric discrete-trait model with Bayesian stochastic search variable selection to reconstruct how clade 2.3.4.4b HPAIV H5 isolated in cattle moved among the eight US states (Idaho, Texas, Kansas, Michigan, New Mexico, North Carolina, Ohio, and South Dakota). The interstate HPAIV movement, inferred by these phylodynamic techniques (Fig. 3), demonstrated that following the first confirmed case in Texas, HPAIV moved rapidly across the US. The phylogenetic signal within the HA gene of cattle B3.13 genes was relatively low, and we conservatively identified when a state transition occurred, i.e., Bayes factor >3 with an associated posterior probability greater than 75%. Using these thresholds, there was phylogenetic evidence for the movement of the B3.13 genotype from Texas to Kansas, Michigan, and New Mexico. There was sufficient evidence in the phylodynamic analyses to support the proposition that the B3.13, following its introduction into Michigan, was moved into North Carolina, but it was more likely that a confirmed Texas to North Carolina animal movement moved the virus as we were unable to confirm a link between Michigan and North Carolina dairy herds. This implies that direct movement of cattle based upon production practices allowed for virus dissemination. The movement of viruses from one region to another will influence the opportunity for reassortment and subsequent changes in genomic diversity. Areas that received large shipments of cattle and their viruses will provide more opportunities for reassortment and the potential emergence of IAV strains with phenotypes that may have increased zoonotic potential.

### Transmission within cattle resulted in low-frequency variants associated with increased transmission efficiency and changes in virulence and pathogenicity

We identified within-host sequence variants across the genome that were present in >0.5% of whole genome sequencing reads, matching to the custom database from the Influenza Research Database (*35*) and a literature search (Table S1). If a sequence variant were to arise that provided a selective advantage, it could increase in frequency through transmission, and potentially alter virus phenotype. Generally, most single-nucleotide variants (SNVs) were present at low frequencies (Data S5). There were 491 amino acid sites with nonsynonymous amino acid changes in cattle; of the variable amino acid sites, 309 nonsynonymous mutations occurred at sites associated with functional changes (mean 44.32 ± 5.06 potential functional changes per sample, range: 24-64). We detected variants associated with changes in pathogenicity in HA, MP, NP, and PB2 (e.g., increase from Q154R in HA and S207G in MP; a decrease in V105M in NP; an increase in D701N in PB2: Table 1: Tables S2-S5 for other animal groups). We also detected SNVs previously associated with changes in virulence (e.g. Q134K/R in HA; increase in V67I in NA; an increase in E627K in PB2) and increased host-range specificity (e.g., A55T, E57G, N71S in N1; and E229K in NS). We detected variants in PB2 associated with mammalian adaptation (an increase in E627K, M631L, and D701N and decrease in V495I), where the frequency was 33% in a single animal for E627K, and at 99% in 214 cattle for M631L. We did not detect the key mammal adaptation mutation PB2 271A in cattle, even at low frequencies; however, one mammal sample contained the mutation in the consensus gene, and we detected mutations on HA that affected receptor binding affinity (Table 1). We calculated synonymous (π*_S_*) and nonsynonymous (π*_N_*) site ratio to assess natural selection; π*_N_/*π*_S_ <*1 suggestive of purifying selection and π*_N_/*π*_S_ >*1 suggestive of positive selection. When we combined the diversity estimates across genes, all cattle strains exhibited π*_N_/*π*_S_ <*1; the gene π*_N_/*π*_S_* estimates varied with none having >1 (HA, π*_N_/*π*_S_* = 0.042; MP, π*_N_/*π*_S_* = 0.566; NA, π*_N_/*π*_S_* = 0.188; NP, π*_N_/*π*_S_* = 0.032; NS, π*_N_/*π*_S_* = 0.355; PA, π*_N_/*π*_S_* = 0.160; PB1, π*_N_/*π*_S_* = 0.048; PB2, π*_N_/*π*_S_* = 0.039), suggesting that the within-host virus populations tended to exhibit weak purifying selection when compared to an ancestral reference. There was no direct evidence that these minor populations of sequence variants alter phenotype of B3.13 genotype HPAIV in cattle. Variants should be monitored to detect whether they increase in frequency as some of the amino acid positions have been associated with increases in pathogenicity, virulence, transmission efficiency and mammalian or human adaptation in laboratory experiments and other animals.

**Table 1.** Sequence variants detected in functionally relevant sites within H5N1 2.3.4.4b strains isolated in cattle that have been associated with pathogenicity, host-adaptation, and virulence. Raw read data from cattle samples were processed and high- and low-frequency single nucleotide variants (SNVs) were identified relative to the most recent common ancestor of the phylogenetic group in this study. The SNVs that induced a coding region change were screened against a database of positions associated with functional change with a relevant selection shown here. The number of cattle samples with the SNV were enumerated, the mean allele frequency was calculated, the presence of the mutation within the consensus gene sequence was determined, and the variants detected at low frequencies were counted.

| Gene | Coding region change | Functional type | Cattle with variant (#) | Mean allele frequency | Consensus sequence | Low frequency variants |
| --- | --- | --- | --- | --- | --- | --- |
| HA | E91K | increase human adapt | 1 | 0.05 | 0 | 1 |
| HA | S137F | change human adapt | 1 | 0.375 | 0 | 1 |
| HA | Q154R | change pathogenicity | 6 | 0.012 | 0 | 6 |
| HA | 172A | increase human adapt | 213 | 0.999 | 213 | 0 |
| HA | A172T | decrease human adapt | 9 | 0.891 | 8 | 1 |
| HA | N209T | change human adapt | 1 | 0.012 | 0 | 1 |
| HA | Q234K/R | change virulence | 9 | 0.044 | 0 | 9 |
| HA | G240R | increase human adapt | 1 | 0.025 | 0 | 1 |
| HA | S336N | change pathogenicity | 17 | 0.885 | 15 | 2 |
| HA | P337L | increase virulence | 8 | 0.715 | 5 | 2 |
| MP | R77K | change pathogenicity | 1 | 0.006 | 0 | 1 |
| MP | S207G | decrease pathogenicity | 4 | 0.013 | 0 | 4 |
| NA | A55T | virulence / host range | 222 | 0.997 | 222 | 0 |
| NA | E57G | virulence / host range | 7 | 0.016 | 0 | 7 |
| NA | V67I | virulence / host range | 222 | 0.997 | 222 | 0 |
| NA | N70S/D | virulence / host range | 221 | 0.99 | 221 | 2 |
| NA | N71S | virulence / host range | 88 | 0.999 | 88 | 0 |
| NA | T438A/I | increase antiviral resist | 3 | 0.627 | 2 | 1 |
| NP | V105M | decrease pathogenicity | 221 | 0.998 | 221 | 0 |
| NS | V205I/G | change virulence | 222 | 0.998 | 222 | 0 |
| NS | E229K | change virulence | 18 | 0.968 | 18 | 0 |
| PB2 | V495I | change mammal adapt | 211 | 0.997 | 211 | 0 |
| PB2 | E627K | increase virulence/adapt | 1 | 0.327 | 0 | 1 |
| PB2 | M631L | increase mammal adapt | 214 | 0.999 | 214 | 0 |
| PB2 | D701N | increase pathogenicity | 2 | 0.015 | 0 | 2 |

## Discussion

The potential for HPAI H5N1 to become endemic in cattle will shape the zoonotic risk of the B3.13 genotype. There may be low levels of immunity against H5N1 viruses (*36–39*) and the immunological landscape in the human population affects disease severity (*40*). Genetically similar viruses do have the potential to cross the species barrier as there has already been a clade 2.3.4.4b B3.13 virus infection in a person with conjunctivitis in March of 2024. The existing prepandemic candidate vaccine viruses (CVV) do retain cross-reactivity with currently circulating clade 2.3.4.4b HPAI H5N1 (*41*). These CVVs are coordinated and shared among the WHO Global Influenza Surveillance and Response Network for use by academic, government, and industry partners for research and development (*42*). However, recent viruses collected in the US had reduced reactivity with the A/Astrakhan/3212/2020 candidate vaccine virus and based on these data and other genetic and epidemiologic measures, a new CVV for the clade 2.3.4.4b viruses was proposed (*41*).

The HPAI H5N1 genotype B3.13 viruses circulating in cattle represent a potential zoonotic threat based on the evidence we present for transmission in a mammalian host. Based upon current information, it appears that once infected, a cow may shed virus for 2-3 weeks. We detected some amino acid mutations at sites associated with mammalian adaptation that had already become fixed in the virus population that likely reflect the ∼4 months of evolution and limited local circulation in dairy cattle. Notably, important low-frequency sequence variants within cattle were also detected, even within the limited time following the first spillover. If these low-frequency variants become dominant, they may have phenotypes that increase the probability of interspecies transmission. Further studies are needed to understand the pathobiology and evolution of the virus in dairy cattle. In addition, there is the potential for multiple animal species to be co-located on agricultural premises, each species may be infected with endemic IAV strains, and an IAV co-infection with HPAI could result in reassortment and the emergence of new strains that increase zoonotic risk (*43, 44*). Monitoring of cattle for HPAI will inform epidemiological risk and provide an early warning for whether this interspecies transmission event and dissemination of the viruses throughout the US dairy cattle herd represents a future threat to human health.

## Materials and Methods

### Sample isolation, whole genome sequencing, assembly, and single nucleotide variant calling

IAV extraction and reverse transcription real-time PCR (RT-rtPCR) were performed at the National Animal Health Laboratory Network member labs and the U.S. Department of Agriculture, National Veterinary Services Laboratories according to the standard operating procedures (*3*). Influenza A virus RNA from samples was amplified (*44*) and after amplification was completed, we generated cDNA libraries for iSeq by using the Illumina DNA Sample Preparation Kit, (M) Tagmentation (Illumina, https://www.illumina.com) and the 300-cycle iSeq Reagent Kit v2 (Illumina) according to manufacturer instructions. We performed reference guided assembly of genome sequences using IRMA v0.6.7 (*45*).

We developed a bioinformatics pipeline for processing, calling single nucleotide variants (SNVs), and analyzing Illumina short read data for influenza A virus called “Flumina” (https://github.com/flu-crew/Flumina). The pipeline uses Python v3.10, R v4.4 (*46*), and SnakeMake (*47*) to organize programs and script execution. A custom Python script organizes raw reads for SnakeMake that subsequently executes other programs for variant calling. The pipeline cleans the raw reads of adapter contamination, low complexity sequences, and other sequencing artifacts using the program FASTP (*48*). IRMA v0.6.7 (*45*) was used on the processed reads to generate consensus contigs that were used for phylogenetic and phylodynamic analyses and other summary statistics and graphs. Following these steps, the pipeline maps reads to a single ancestral reference strain, which was generated from ancestrally reconstructed sequences as described below, by indexing using BWA (*bwa index -a bwtsw* function; (*49*)). Then it uses SAMtools (*50*) to create an fai index with the *faidx* function and generate a sequence dictionary for GATK v4.4 (*51*) using the *CreateSequenceDictionary* function.

For calling of high frequency SNVs, we used GATK v4.4 (*51*) following best practices for discovering and calling SNVs (*52*). The pipeline processes the cleaned reads through the GATK4 functions FastqToSam, RevertSam, and AddorReplaceReadGroups. Next, the BAM of the reads are converted back to Fastq and mapped with BWA using the function *bwa mem -M*. The mapped reads BAM file are merged with the unmapped reads BAM, and duplicate reads are marked after sorting the BAM file by coordinate with the functions *SortSam* and *MarkDuplicates*. Finally, *HaplotypeCaller* is used (parameters: -ERC GVCF -ploidy 1) to call haplotypes, and *GenotypeGVCFs* is used to genotype the sample. Specific variants were selected using the *SelectVariants* function for the SNP type of variant. Using *VariantFiltration*, a filtered set of variants were created by applying the following filters: Qual < 30 (Quality), QD < 2 (Quality by Depth), SOR < 3 (StrandOddsRatio), FS < 60 (FisherStrand), MQ < 40 (MapQuality), MQRankSum < 12.5 (MapQuality RankSumTest), and ReadPosRankSum < 8 (ReadPosRankSumTest). To call low frequency SNVs, the program LoFreq (*53*) was applied, with the similar preprocessing steps where the same BAM was used as input from *HaplotypeCaller*. A database was generated using the Sequence Feature Variant Types tool from the Influenza Research Database (*34*) for all eight genes, and SNVs associated with nonsynonymous changes in each gene were screened to determine whether these had previously been associated with phenotypic change. To estimate genome-wide estimates of natural selection, we used the program SNPGenie on the VCF files (*54*).

### Data sources, curation, and preparation

To generate a reference dataset of relevant HPAI H5N1 strains, we queried GISAID (*55*) for all H5 genomes with 8 complete gene segments available collected between the dates of January 01 2020 and March 29 2024 in the United States. This query provided 23,322 sequences from 2,915 genomes: we extracted the hemagglutinin (HA) gene sequence and aligned the data using MAFFT v7.525 (*56*) with default parameters. We subsequently inferred a maximum likelihood phylogenetic tree with IQ-Tree v2.2.0 (*57*) following automatic model selection (*58*). Based on this tree, we classified all data using a custom python clade identifier called GenoFLU (https://github.com/USDA-VS/GenoFLU) and filtered the data to only those strains from the HPAI clade 2.3.4.4b. We then removed identical HA sequences, ensuring that we retained the cattle strains that had been published in GISAID and the associated human case (A/Texas/37/2024, GISAID accession EPI_ISL_19027114). This process resulted in a reference gene dataset of 1393 strains. For Bayesian phylodynamic analyses, these data were then subsampled using smot v.1.0.0 (*59*) where we maintained representation by sampling 20% of the tips within each monophyletic clade on the HA gene phylogeny inferred using FastTree v2.1.11 (*60*). Using the subsampled dataset of n=964 strains, we inferred the ancestral sequences of North American 2.3.4.4b strains for each gene segment using TreeTime v0.9.4 ancestral reconstruction functionality. The ancestral sequences were reconstructed at the deepest nodes on each gene tree that comprised at least 99% of all strains in the subsampled dataset.

### Phylogenetic analyses

The subsampled reference data were merged with whole genome sequences generated within this study. For each gene segment, the data were aligned with MAFFT v7.525 (*56*) with default settings, and maximum likelihood (ML) trees were inferred with IQ-Tree v2.2.0 (*57*) following automatic model selection (*58*). These trees were used to assess root-to-tip divergence using TempEst v1.5.3 (*61*) and genes with incongruent divergence and sampling date were removed. Subsequently, we realigned each gene dataset, and inferred new ML phylogenetic trees using IQ-Tree v2.2.0. Using the ML phylogenies for each gene segment as a starting point, we generated time-scaled phylogenies, rooted under the strict molecular clock assumption using the TreeTime v0.8.4 phylodynamic toolkit (*62*). TreeTime was executed with the GTR site substitution model with 10 iterations of optimization. The host state (poultry, wild bird, cattle, mammal, domestic mammal, and humans) and the US location were inferred for each ancestral node on the time-scaled phylogeny using a TreeTime migration model. For each ancestral node, TreeTime inferred a probability of the respective virus belonging to the specific discrete trait (i.e., Texas or New Mexico; wild bird or cattle) and we subsequently assigned the most likely trait (over 50% probability) to each node. This approach allowed us to identify branches on the tree where spillovers may have occurred and between different locations and this approach complemented our Bayesian phylodynamic analyses.

We also tracked the emergence of single nucleotide variants (SNVs) within clade 2.3.4.4b consensus sequences that were generated using IRMA v0.6.7 (*45*). SNVs were annotated and compared using a custom pipeline called vSNP3 (https://github.com/USDA-VS/vSNP). The pipeline infers phylogenetic trees using RAxML (*63*) and generates tables of SNVs relative to a reference composed of two North American wild bird origin segments (PB2, NP) and six segments from a H5N1 clade 2.3.4.4b clade virus. These data were used to determine viral genome sequence similarity and to identify genomic links between premises (Fig. 1, Fig. S1, Data S2).

### Reassortment analyses

We assembled a separate dataset for reassortment analysis that comprehensively included all H5N1 whole genomes collected from 2020 to present through downloading n=27,177 HPAI H5N1 gene sequences from GISAID [accessed April 19, 2024] (*55*). We maintained only those strains with complete sequences for each of the eight gene segments, resulting in n=2893 whole genomes. We merged these data with the assembled whole genomes from our study and generated alignments for each gene using MAFFT v7.490 (*56*) with default parameters. We subsequently inferred phylogenetic trees for each gene segment using IQ-Tree v1.6.12 (*64*) under the generalized time-reversible (GTR) model of nucleotide substitutions with the stationary probabilities estimated from the empirical base frequencies and 5 free-rate categories (*65*). To identify reassortment events on the HA phylogeny, we applied TreeSort v.0.1.1 (https://github.com/flu-crew/TreeSort) with a maximum molecular clock deviation parameter of 2.5. TreeSort uses TreeTime (*62*) to estimate the substitution rates for each gene segment and identifies branches on a tree with a high signal of reassortment. We then removed n=35 strains that were molecular clock outliers in the HA gene and were not related to the outbreak in dairy cattle and repeated the TreeSort analysis.

### Phylogenomic analyses over concatenated B3.13 genomes

To improve the phylogenetic signal among the cattle and associated sequences, we concatenated the coding regions of all gene segments into whole genome sequences for all reassortment-free B3.13 strains from the reassortment analysis above. This resulted in n=256 aligned whole genome sequences, including n=220 cattle strains. We inferred a phylogenetic tree from these sequences using an edge-linked partition model in IQ-Tree v.2.3.2 (*57*), where every partition corresponded to an individual gene segment and was associated with a different GTR model with empirical base frequencies and 5 free-rate categories. IQ-Tree was run with ultrafast bootstrap with 1000 replicates (*66*). We then used the resulting tree topology as input to TreeTime v.0.9.4 (*62*) to infer a time-scaled tree of the B3.13 genomes. TreeTime was executed with an option to account for covariation in tip dates due to shared ancestry.

### Inference of B3.13 transmission chains in dairy cattle

Inference of transmission chains between hosts, sampled diversity, sampling time, and generation time were estimated from the temporally scaled genome tree using the TransPhylo R package (*67, 68*). As input, we used the B3.13 tree generated in the phylogenomic analysis. Multifurcations in the tree were suppressed to bifurcations and all edge weights were rounded to at least 1e-9. The prior generation time and sampling density were gamma-distributed parameters extracted from published studies (75, 76) resulting in a mean of 5 days (*69, 70*) and a variance of 2 days. We adapted protocol 3 (*68*) ensuring an Effective Sample Size (ESS) > 200 at the end of the simulations. The MCMC simulations were run for 1e7 iterations, discarding the first 5% as burn-in. After the simulations converged, the medoid tree from the posterior set of inferred trees was used as the final inferred transmission tree. Transmissions of HPAIV from host to host (e.g., wild bird to cattle) were inferred by TransPhylo (75, 76) and the distribution of incident cases (sampled and unsampled) was generated by adapting protocol 4 (Fig. S8) (*68*).

### Bayesian evolutionary inference and discrete phylogeography

We estimated when HPAI H5N1 clade 2.3.4.4b were introduced into dairy cattle under a Bayesian framework in BEAST v.1.10.4 (*71*) with the BEAGLE library (*72, 73*). We merged the subsampled public data with newly generated whole genome data and aligned the hemagglutinin dataset (n=606 HA genes). We implemented a generalized time reversible (GTR) nucleotide substitution model (*74*) with gamma-distributed site heterogeneity (*75*), an uncorrelated relaxed clock with lognormal distribution (*76*), and a Gaussian Markov random field (GMRF) Bayesian skyride with time aware-smoothing for the coalescent model (*77*). We conducted six independent Markov chain Monte Carlo (MCMC) sampling runs with 100 million iterations with sampling every 10,000 iterations. The results were analyzed using the GMRF skyride reconstruction in Tracer v1.7.2 (*78*), and runs were combined to ensure an effective sample size of more than 200. A time-scaled maximum clade credibility (MCC) tree was generated using TreeAnnotator v1.8.4 using median node heights and 20 percent burn-in (*71*). Dates of time to the most recent common ancestor (TMRCA) were inferred from the nodes between clades, using the 95 percent higher posterior density (HPD) as the range of uncertainty.

To reconstruct the spatial diffusion of the H5N1 virus across states within the US and different hosts (poultry, wild bird, cattle, mammal, domestic mammal, and humans), we applied an asymmetric discrete-trait phylogeographic analyses with Bayesian stochastic search variable selection, a strict molecular clock, and an exponential growth model in BEAST v.1.10.4 (*71*). We conducted this analysis on the subsampled and aligned hemagglutinin dataset and an extracted the monophyletic clade of cattle HA clade 2.3.4.4b genes with any host category (host) or cattle only (location). In this approach, each HA gene had a US state and host group designated as a trait, and transitions from one category to another (e.g. from Texas to New Mexico) was inferred along the internal branches representing the evolutionary history of the virus. These transitions are termed Markov jumps and state change counts for the HA clade trait were reconstructed in BEAST v1.10.4. We conducted at least 5 independent Markov chain Monte Carlo (MCMC) chains with 50 million iterations with sampling every 5,000 iterations. Convergence and mixing was assessed in Tracer v1.7.2, as described above, and runs were combined to ensure an effective sampling size of more than 200. We used the resulting posterior trees to provide estimates of the ancestral region and host for each internal node. We then used SpreaD3 v.0.9.6 (*79*) to estimate the Bayes Factor (BF) from the estimated state transition rates and used these values as statistical support for spatial movements: we imposed thresholds used in previously published studies, i.e., a definitive spatial movement had a BF > 100, sufficient evidence was assessed as 100 > BF > 3 (*1, 80*).

## Acknowledgments

We gratefully acknowledge producers, veterinarians, and laboratories for participating in the epidemiological investigations and sharing the results of sequence analysis. We thank the U.S. Department of Agriculture (USDA), Animal and Plant Health Inspection Services veterinary epidemiology team for contact tracing, sample collection, and coordinating the response to the outbreak: Nicole Amey, Bradley Christenson, Michael Contente, Lindsey Holmstrom, Dillon McBride, Stacey Schwabenlander, and Jason Lombard (Colorado State University). We thank Michael Zeller from Iowa State University for insight into TransPhylo, and Brian Stucky and Jeffrey Silverstein for providing priority access to the SCInet High Performance Computing infrastructure at the U.S. Department of Agriculture, Agricultural Research Service. We also acknowledge all data contributors, i.e., the Authors and their Originating laboratories responsible for obtaining the specimens, and their submitting laboratories for generating the genetic sequence and metadata and sharing via the GISAID Initiative, on which components of this research was based. Mention of trade names or commercial products in this article is solely for the purpose of providing specific information and does not imply recommendation or endorsement by the U.S. Department of Agriculture (USDA), DOE, or ORISE. The funders had no role in study design, data collection and interpretation, or the decision to submit the work for publication. The findings and conclusions in this publication are those of the authors and should not be construed to represent any official USDA or U.S. Government determination or policy. USDA is an equal opportunity provider and employer.

## Funding

National Institute of Allergy and Infectious Diseases, National Institutes of Health, Department of Health and Human Services contract 75N93021C00015 (TKA, ALB)

U.S. Department of Agriculture Agricultural Research Service project 5030-32000-231-000-D (ALB, TKA)

U.S. Department of Agriculture Agricultural Research Service project 0201-88888-003-000D (ALB, TKA)

U.S. Department of Agriculture Agricultural Research Service project 0201-88888-002-000D (ALB, TKA)

U.S. Department of Agriculture Agricultural Research Service Research Participation Program of the Oak Ridge Institute for Science and Education through the U.S. Department of Energy contract DE-SC0014664 (GJ, CH, TN, SW)

Centers for Disease Control and Prevention of the U.S. Department of Health and Human Services Interagency Agreement DE-SC0000001 (ALB, TKA)

## Author contributions

Conceptualization: MKT, TKA

Data curation: TN, CH, BI, KL, MLK

Formal Analysis: TN, CH, AM, MT, KL, SV, SW, GMJ, BI, SML, MKT, TKA

Funding acquisition: ALB, SRA, MKT, TKA

Investigation: KL, MLK, SML, SCA, KRJ

Methodology: TN, CH, AM, MT, KL, MLK, GMJ, SV, SW, BI

Project administration: TKA, MKT

Resources: DRM, GL, DGD, EAF, KMD, AKS, ACT, KRS, DLS, ES, SCA, KRJ, SRA

Software: CH, AM, SML

Supervision: MKT, TKA

Visualizations: TN, AM, GMJ, SML, SV, TKA

Writing – original draft: CH, AM, MKT, TKA

Writing – review & editing: TN, CH, AM, MT, KL, MLK, GMJ, SV, SW, DRM, GL, DGD, EF, KMD, AKS, ACT, KRS, DLS, ES, SML, SCA, ALB, SRA, MKT, TKA

## Data and materials availability

All data, code, and materials used in the analysis are available at https://github.com/flu-crew/dairy-cattle-hpai-2024. Sequence data generated within this study are provided at NCBI GenBank and NCBI SRA and the accession numbers are provided in the supplementary text data files.

## Supplementary Text

Figs. S1 to S9

Tables S1 to S5

References (*45–81*)

Data S1 to S5

