## Supplementary Material for "Emergence and interstate spread of highly pathogenic avian influenza A(H5N1) in dairy cattle"

**The PDF file includes:**

Figs. S1 to S9

Tables S1 to S5

References (45-81)

**Other Supplementary Materials for this manuscript include the following:**

Data S1 to S5

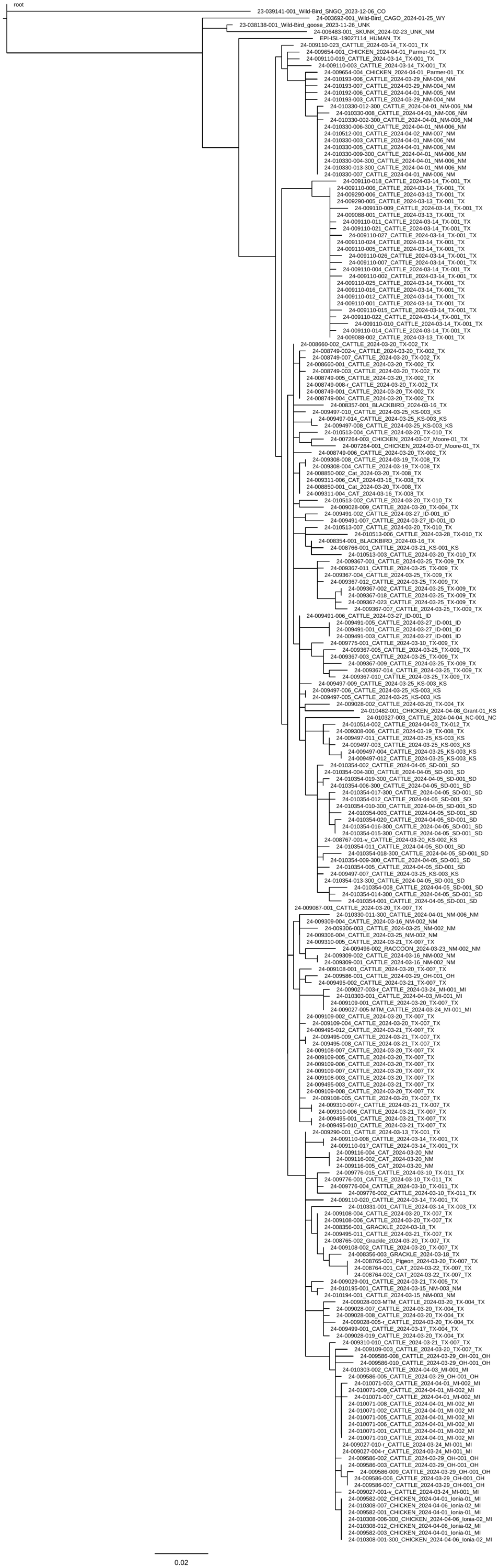

Fig. S1.

Phylogenetic tree inferred from single nucleotide variants (SNVs) within clade 2.3.4.4b consensus sequences. SNVs were annotated and compared using a custom pipeline called vSNP3 (<https://github.com/USDA-VS/vSNP>). The pipeline infers phylogenetic trees using RAxML (*63*) and generates tables of SNVs relative to a reference composed of two North American wild bird origin segments (PB2, NP) and six segments from a H5N1 clade 2.3.4.4b clade virus. These data were used to determine viral genome sequence similarity and to identify genomic links between premises (Fig. 1).

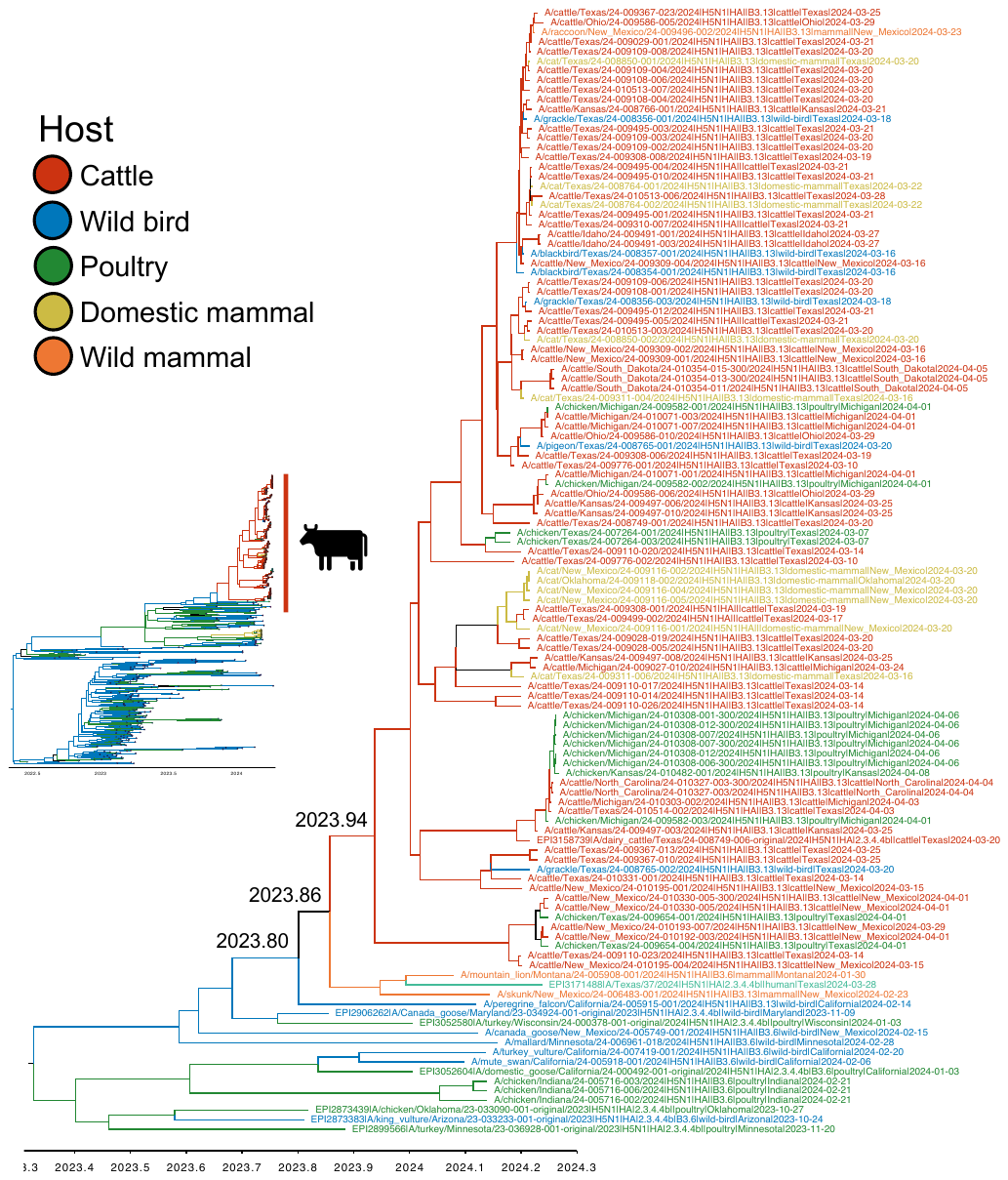

Fig. S2.

A paraphyletically subsampled section of the full time-scaled phylogeny of the HA gene between December 2021 and April 2024 demonstrating the single introduction of the virus from wild birds to dairy cattle with an estimated date of December 2023 (95% credible interval: October 2023 – January 2024). B3.13 strains inherited PB1, PA, HA, NA, MP, and NS genes from a B3.6 ancestor and acquired different PB2 and NP genes from North American LPAI viruses. This phylogenetic tree topology demonstrates approximately 12 subsequent spillovers from cattle to domestic cats, poultry, and peridomestic animals.

Fig. S3.

Time-scaled maximum-likelihood trees of HPAI H5N1 sequences in the 2.3.4.4b-clade HA and NA segments (n=601). Tips of taxa in the cattle clade is labelled in red. The time scale for the HA tree is 2023-2024.5 and the scale of the NA tree is 2022.5-2024.5. These trees are also hosted in a git repo <https://github.com/flu-crew/dairy-cattle-hpai-2024> as nexus tree files and as searchable PDFs.

Fig. S4.

Time-scaled maximum-likelihood trees of HPAI H5N1 sequences in the 2.3.4.4b-clade MP and NP segments (n=601). Tips of taxa in the cattle clade is labelled in red. The time scale for the MP tree is 2022.5-2024.5 and the scale of the NP tree is 2021-2025. These trees are also hosted in a git repo <https://github.com/flu-crew/dairy-cattle-hpai-2024> as nexus tree files and as searchable PDFs.

Fig. S5.

Time-scaled maximum-likelihood trees of HPAI H5N1 sequences in the 2.3.4.4b-clade NS and PA segments (n=601). Tips of taxa in the cattle clade is labelled in red. Tips of taxa in the cattle clade is labelled in red. The time scale for the NS tree is 2023-2024.5 and the scale of the PA tree is 2010-2025. These trees are also hosted in a git repo <https://github.com/flu-crew/dairy-cattle-hpai-2024> as nexus tree files and as searchable PDFs.

Fig. S6.

Time-scaled maximum-likelihood trees of HPAI H5N1 sequences in the 2.3.4.4b-clade PB1 and PB2 segments (n=601). Tips of taxa in the cattle clade is labelled in red. Tips of taxa in the cattle clade is labelled in red. The time scale for the PB1 tree is 2010-2025 and the scale of the PB2 tree is 2023.4-2024.3. These trees are also hosted in a git repo <https://github.com/flu-crew/dairy-cattle-hpai-2024> as nexus tree files and as searchable PDFs.

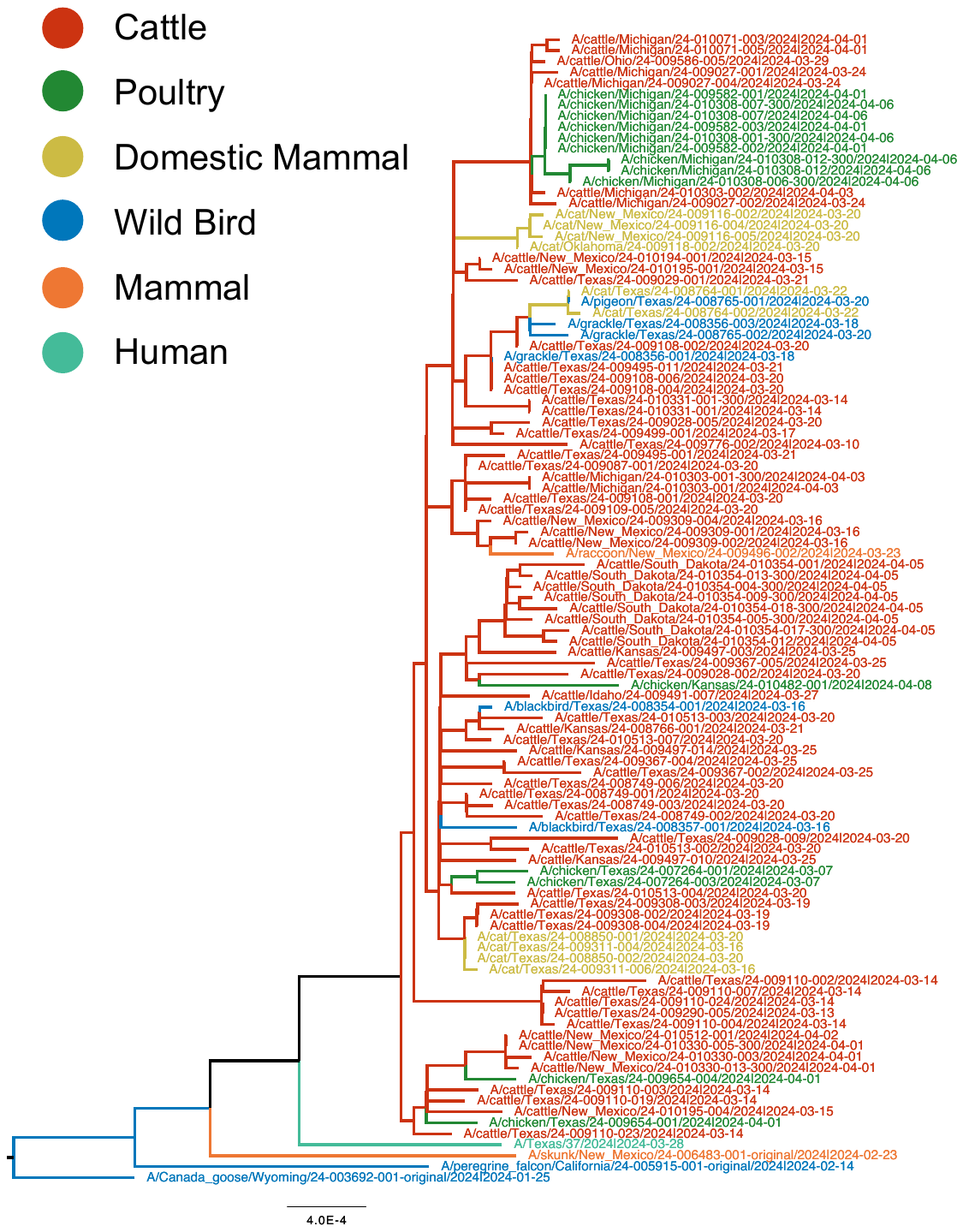

**Fig. S7.**

A hypothesis on the evolutionary history of the H5N1 2.3.4.4b B3.13 genotype that emerged in cattle and a range of other mammalian and avian hosts. This phylogeny was inferred using maximum likelihood methods from concatenated genomes: this data indicated approximately 12 cattle to other host transmission events. The phylogeny was paraphyletically subsampled for visualization purposes.

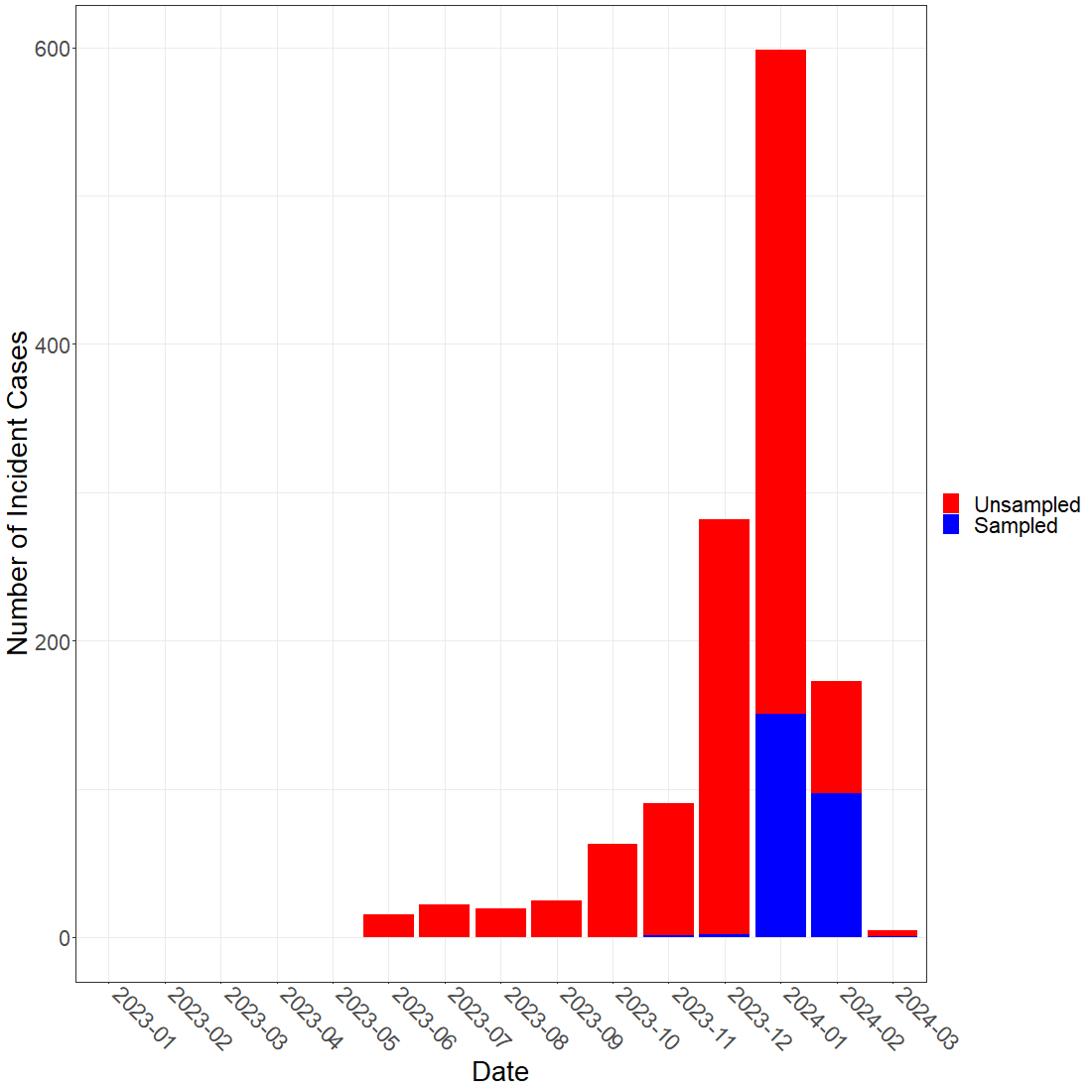

**Fig. S8.**

Inference of sampled and unsampled transmission chains through a TransPhylo reconstruction of the H5N1 clade 2.3.4.4b B3.13 genotype outbreak in dairy cattle. These data represent inferred numbers of sampled and unsampled cases over time in the posterior transmission trees produced using TransPhylo.

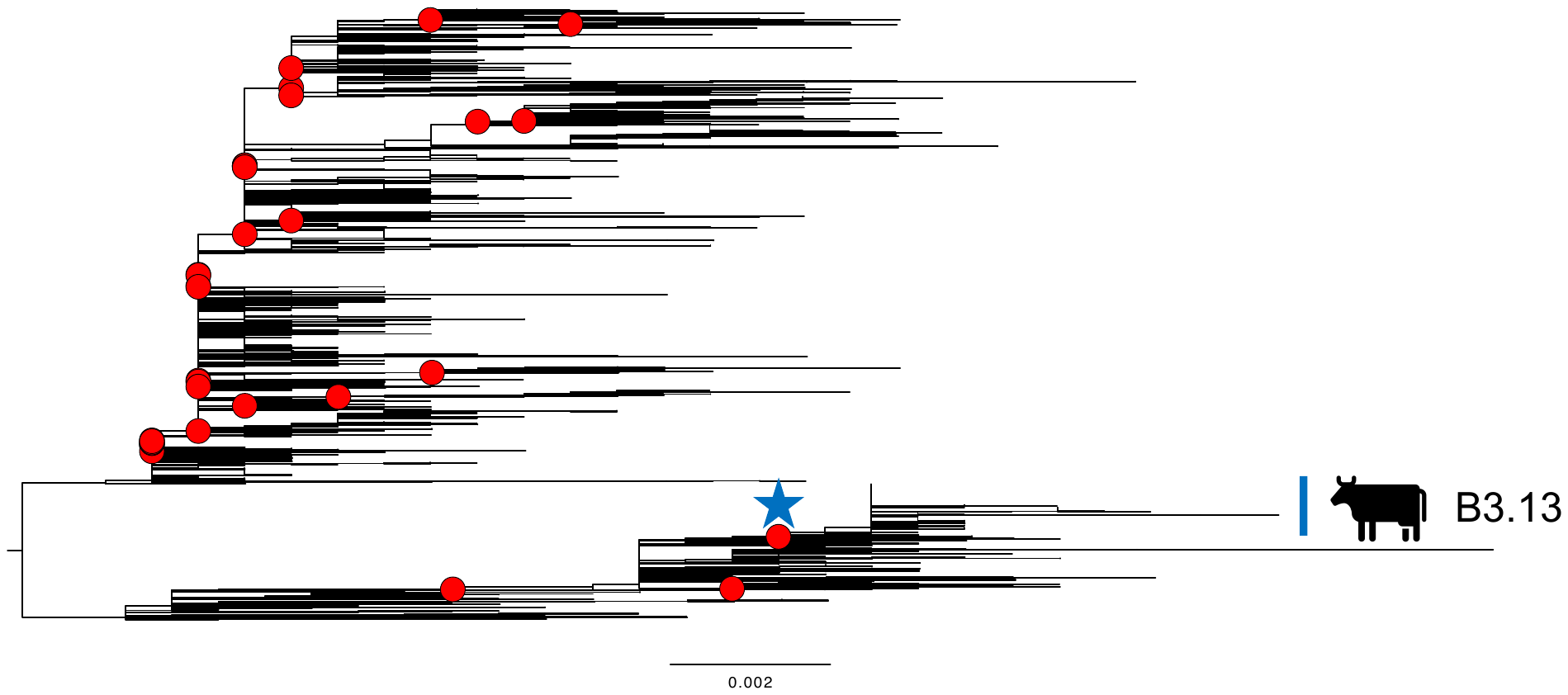

Fig. S9.

Reassortment events across the 2.3.4.4b HA phylogeny. Red circles represent 24 major reassortment events with more than 20 downstream detections. The blue star highlights the reassortment event that led to the emergence of the B3.13 genotype in dairy cattle and a range of other mammalian and avian hosts. These trees are also hosted in a git repo <https://github.com/flu-crew/dairy-cattle-hpai-2024> as nexus tree files and as searchable PDFs.

Table S1.

**Table S1. Functionally relevant sequence variants screened within H5N1 strains isolated in various animals from this study that have been associated with pathogenicity, host-adaptation, and virulence.** A database was generated using the Sequence Feature Variant Types tool from the Influenza Research Database and reduced to include functional annotations with the keywords: pathogenicity, host-adaptation, virulence, transmission, host-range expansion and species adaptation. Additional functional variants were added from a literature search to include those not in the database (marked with a *). Presented here are the select functional changes show in Table 1 and reported in the main text with the NCBI PubMed identified associated with the study.

| Gene | Amino acid position | Short description | Citation |
| --- | --- | --- | --- |
| HA | 91 | binding affinity to human-type receptor | PMID:17108965 |
| HA | 137 | binding affinity to human-type receptor | PMID:20427525 |
| HA | 154 | pathogenicity | PMID: 15331729 |
| HA | 158 | binding affinity to mammal receptor, transmission | PMID: 20041223* |
| HA | 160 | binding affinity to human-type receptor | PMID: 17108965 |
| HA | 172 | binding affinity to human-type receptor | PMID:18672252, 20427525 |
| HA | 182 | binding affinity to human-type receptor | PMID: 17108965; 20392847 |
| HA | 192 | binding affinity to human-type receptor | PMID: 17108965, 21637809* |
| HA | 209 | binding affinity to human-type receptor | PMID:17108965 |
| HA | 228 | binding affinity to human-type receptor | PMID: 15331729 |
| HA | 234 | increases replication efficiency of virus | PMID:20519408, 18632950 |
| HA | 240 | binding affinity to mammal receptor | PMID:20392847, 22056389 |
| HA | 336 | pathogenicity | PMID: 20881092 |
| HA | 337 | virulence in mammals | PMID: 22278228 |
| MP | 30 | virulence in mammals | PMID:19117585 |
| MP | 43 | drug site binding | PMID: 18235503* |
| MP | 77 | virulence | PMID: 16699003 |
| MP | 139 | virulence in mammals | PMID:8879138, 10426210 |
| MP | 207 | virulence | PMID:23209789 |
| MP | 215 | virulence | PMID:19117585 |
| NA | 55 | virulence / host range | PMID: 19609439 |
| NA | 57 | virulence / host range | PMID: 19609439, 16525739 |
| NA | 67 | virulence / host range | PMID: 19609439 |
| NA | 70 | virulence / host range | PMID: 8419645 |
| NA | 71 | virulence / host range | PMID: 8419645 |
| NA | 110 | viral replication and severity | PMID: 28597818* |
| NA | 295 | drug site binding | PMID: 37494978* |
| NA | 438 | drug site binding | PMID: 37494978* |
| NA | 453 | viral replication and severity | PMID: 28597818* |
| NP | 105 | pathogenicity | PMID: 21123376 |
| NP | 319 | polymerase activity in mammal cells | PMID: 16339318 |
| NS | 42 | binding affinity to human-type receptor | PMID: 18032512 |
| NS | 91 | region associated with pathogenicity | PMID: 20854176 |
| NS | 92 | region associated with pathogenicity | PMID: 20854176 |
| NS | 93 | region associated with pathogenicity | PMID: 20854176 |
| NS | 94 | region associated with pathogenicity | PMID: 20854176 |
| NS | 95 | region associated with pathogenicity | PMID: 20854176 |
| NS | 99 | virulence and pathogenicity | PMID: 20854176 |
| NS | 101 | virulence | PMID: 10873787 |
| NS | 103 | binding affinity to human-type receptor | PMID: 19052083; 21593152 |
| NS | 106 | binding affinity to human-type receptor | PMID: 19052083; 21593152 |
| NS | 125 | binding affinity to mammal receptor, pathogenicity | PMID: 18983930 |
| NS | 205 | virulence | PMID: 20862325 |
| NS | 229 | virulence | PMID: 18334632 |
| PA | 37 | polymerase activity in mammal cells | PMID: 24371069* |
| PA | 383 | polymerase activity in mammal cells | PMID: 18615018 |
| PA | 409 | polymerase activity in mammal cells | PMID: 18615018 |
| PB1 | 3 | polymerase activity in mammal cells | PMID:16533883 |
| PB1 | 622 | virulence | PMID: 26656683* |
| PB2 | 158 | viral replication and severity | PMID: 28597818* |
| PB2 | 271 | polymerase activity in mammal cells | PMID: 20181719 |
| PB2 | 389 | polymerase activity in mammal cells | PMID: 27889648* |
| PB2 | 495 | mammal polymerase activity | PMID:19393699 |
| PB2 | 591 | viral replication | PMID: 20700447 |
| PB2 | 598 | polymerase activity in mammal cells | PMID: 27889648* |
| PB2 | 627 | virulence, adapt to mammals | PMID: 19119420 |
| PB2 | 631 | viral replication and severity in mammals | PMID: 28597818* |
| PB2 | 701 | transmission in mammals | PMID: 19119420 |

Table S2.

**Table S2.** **Sequence variants detected in functionally relevant sites within H5N1 2.3.4.4b strains isolated in wild birds that have been associated with pathogenicity, host-adaptation, and virulence.** Raw read data from cattle samples were processed and high- and low-frequency single nucleotide variants (SNVs) were identified. The SNVs that induced a coding region change were screened against a database of positions associated with functional change with a relevant selection shown here. The number of wild bird samples with the SNV were enumerated, the mean allele frequency was calculated, the presence of the mutation within the consensus gene sequence was determined, and the variants detected at low frequencies were counted.

| Gene | Coding region change | Functional type | | Animals with variant (#) | | Mean allele frequency | Consensus sequence | Low frequency variants |
| --- | --- | --- | --- | --- | --- | --- | --- | --- |
| HA | A172E | | pathogenicity | 4 | 0.751 | | 3 | 1 |
| HA | E91K | | human adapt | 1 | 0.032 | | 0 | 1 |
| HA | Q154R | | pathogenicity | 2 | 0.999 | | 2 | 0 |
| HA | Q234K | | virulence | 5 | 0.244 | | 1 | 4 |
| HA | S336N | | pathogenicity | 6 | 0.933 | | 6 | 0 |
| MP | A215V/E | | virulence | 3 | 0.008 | | 0 | 3 |
| MP | S207N/G | | virulence | 2 | 0.011 | | 0 | 2 |
| MP | T139I | | virulence | 1 | 0.022 | | 0 | 1 |
| NA | A55T | | virulence | 167 | 0.997 | | 167 | 0 |
| NA | E57K | | virulence | 2 | 0.999 | | 2 | 0 |
| NA | N295D | | antiviral resistance | 1 | 0.006 | | 0 | 1 |
| NA | N70S | | host range | 164 | 0.998 | | 164 | 0 |
| NA | N71S | | host adaptation | 2 | 0.999 | | 2 | 0 |
| NA | T438I | | antiviral resistance | 4 | 0.51 | | 2 | 2 |
| NA | V453M/G | | mammal adaptation | 13 | 0.46 | | 6 | 7 |
| NA | V67I | | virulence | 8 | 0.998 | | 8 | 0 |
| NP | N319K | | mammal adaptation | 9 | 0.673 | | 6 | 3 |
| NP | V105M | | pathogenicity | 27 | 0.925 | | 25 | 2 |
| NS | D125G | | pathogenicity | 1 | 0.085 | | 0 | 1 |
| NS | E229* | | virulence | 1 | 0.04 | | 0 | 1 |
| NS | V205I/G/D | | pathogenicity | 167 | 0.995 | | 167 | 1 |
| PA | D383E | | mammal adaptation | 1 | 0.102 | | 0 | 1 |
| PA | S409G | | mammal adaptation | 2 | 0.998 | | 2 | 0 |
| PB2 | D701G/Y/E | | host adaptation | 5 | 0.029 | | 0 | 5 |
| PB2 | M631V/L | | mammal adaptation | 9 | 0.89 | | 8 | 1 |
| PB2 | Q591K/H/* | | host adaptation | 3 | 0.382 | | 1 | 2 |
| PB2 | R389K | | mammal adaptation | 3 | 0.609 | | 2 | 1 |
| PB2 | T271A | | mammal adaptation | 1 | 0.022 | | 0 | 1 |
| PB2 | V495I | | mammal adaptation | 8 | 0.876 | | 7 | 1 |

Table S3.

**Table S3.** **Sequence variants detected in functionally relevant sites within H5N1 2.3.4.4b strains isolated in poultry that have been associated with pathogenicity, host-adaptation, and virulence.** Raw read data from cattle samples were processed and high- and low-frequency single nucleotide variants (SNVs) were identified. The SNVs that induced a coding region change were screened against a database of positions associated with functional change with a relevant selection shown here. The number of poultry samples with the SNV were enumerated, the mean allele frequency was calculated, the presence of the mutation within the consensus gene sequence was determined, and the variants detected at low frequencies were counted.

| Gene | Coding region change | Functional type | Animals with variant (#) | Mean allele frequency | Consensus sequence | Low frequency variants |
| --- | --- | --- | --- | --- | --- | --- |
| HA | Q154L | pathogenicity | 2 | 0.996 | 2 | 0 |
| HA | S336N | pathogenicity | 3 | 0.996 | 3 | 0 |
| HA | W192C | host adaptation | 1 | 0.018 | 0 | 1 |
| NA | A55T | increase virulence | 51 | 0.993 | 51 | 0 |
| NA | E57G | virulence | 2 | 0.01 | 0 | 2 |
| NA | N295S | antiviral resistance | 1 | 0.22 | 0 | 1 |
| NA | N70S | host range | 51 | 0.997 | 51 | 0 |
| NA | N71S | host adaptation | 3 | 0.998 | 3 | 0 |
| NA | S110F | mammal adaptation | 2 | 0.046 | 0 | 2 |
| NA | T438A/I | antiviral resistance | 3 | 0.414 | 1 | 2 |
| NA | V453L/G | mammal adaptation | 2 | 0.029 | 0 | 2 |
| NA | V67A/I | virulence | 21 | 0.957 | 20 | 1 |
| NP | V105M | pathogenicity | 23 | 0.975 | 23 | 0 |
| NS | E229K | virulence | 1 | 0.093 | 0 | 1 |
| NS | V205I/G | pathogenicity | 51 | 0.998 | 51 | 0 |
| PA | D383N | mammal adaptation | 1 | 0.017 | 0 | 1 |
| PB2 | D701H | host adaptation | 1 | 0.014 | 0 | 1 |
| PB2 | E627K | virulence | 8 | 0.995 | 8 | 0 |
| PB2 | M631I/L | mammal adaptation | 22 | 0.926 | 20 | 2 |
| PB2 | V495I | mammal adaptation | 20 | 0.998 | 20 | 0 |

Table S4.

**Table S4.** **Sequence variants detected in functionally relevant sites within H5N1 2.3.4.4b strains isolated in mammals that have been associated with pathogenicity, host-adaptation, and virulence.** Raw read data from mammal samples were processed and high- and low-frequency single nucleotide variants (SNVs) were identified. The SNVs that induced a coding region change were screened against a database of positions associated with functional change with a relevant selection shown here. The number of mammal samples with the SNV were enumerated, the mean allele frequency was calculated, the presence of the mutation within the consensus gene sequence was determined, and the variants detected at low frequencies were counted.

| Gene | Coding region change | Functional type | Animals with variant (#) | Mean allele frequency | Consensus sequence | Low frequency variants |
| --- | --- | --- | --- | --- | --- | --- |
| NA | A55T | virulence | 14 | 0.981 | 14 | 0 |
| NA | N70S | host range | 15 | 0.999 | 15 | 0 |
| NA | V67I | virulence | 3 | 0.997 | 3 | 0 |
| NP | V105M | pathogenicity | 2 | 0.996 | 2 | 0 |
| NS | V205G/I | pathogenicity | 15 | 0.997 | 15 | 0 |
| PB2 | D701N | host adaptation | 2 | 0.134 | 0 | 2 |
| PB2 | E627K | virulence | 6 | 1 | 6 | 0 |
| PB2 | M631L | mammal adaptation | 1 | 1 | 1 | 0 |
| PB2 | T271A | mammal adaptation | 1 | 0.997 | 1 | 0 |
| PB2 | V495I | mammal adaptation | 2 | 0.993 | 2 | 0 |

Table S5.

**Table S5.** **Sequence variants detected in functionally relevant sites within H5N1 2.3.4.4b strains isolated in domestic mammals that have been associated with pathogenicity, host-adaptation, and virulence.** Raw read data from cattle samples were processed and high- and low-frequency single nucleotide variants (SNVs) were identified. The SNVs that induced a coding region change were screened against a database of positions associated with functional change with a relevant selection shown here. The number of domestic mammal samples with the SNV were enumerated, the mean allele frequency was calculated, the presence of the mutation within the consensus gene sequence was determined, and the variants detected at low frequencies were counted.

| Gene | Coding region change | Functional type | Animals with variant (#) | Mean allele frequency | Consensus sequence | Low frequency variants |
| --- | --- | --- | --- | --- | --- | --- |
| MP | S207N | virulence | 1 | 0.01 | 0 | 1 |
| NA | A55T | virulence | 27 | 0.995 | 27 | 0 |
| NA | V67I | virulence | 11 | 0.997 | 11 | 0 |
| NA | N70S | host range | 27 | 0.999 | 27 | 0 |
| NA | N71S | host adaptation | 4 | 0.998 | 4 | 0 |
| NP | V105M | pathogenicity | 11 | 0.998 | 11 | 0 |
| NS | V205G/I | pathogenicity | 27 | 0.998 | 27 | 0 |
| PB2 | T271A | mammal adaptation | 1 | 0.999 | 1 | 0 |
| PB2 | V495I | mammal adaptation | 10 | 0.995 | 10 | 0 |
| PB2 | Q591K | host adaptation | 3 | 0.064 | 0 | 3 |
| PB2 | M631L | mammal adaptation | 11 | 0.997 | 11 | 0 |

Data S1.

H5N1 2.3.4.4b strains associated with the dairy cattle outbreak, that were subsequently sequenced, with raw sequence reads archived at the NCBI Sequence Read Archive.

Data S2.

Single nucleotide variant (SNVs) analysis within clade 2.3.4.4b consensus sequences. SNVs were annotated and compared using a custom pipeline called vSNP3 (<https://github.com/USDA-VS/vSNP>). The pipeline infers phylogenetic trees using RAxML (*63*) and generates tables of SNVs relative to a reference composed of two North American wild bird origin segments (PB2, NP) and six segments from a H5N1 clade 2.3.4.4b clade virus. These data were used to determine viral genome sequence similarity and to identify genomic links between premises.

Data S3.

GISAID acknowledgment metadata file.

Data S4.

NCBI Genbank Accession numbers for strains generated in this study.

Data S5.

Data table of all the variants matching to functional sites detected within within clade 2.3.4.4b raw sequence reads.
